# Reversible Sandwich-Based Particle Nanoswitch for Continuous Protein Monitoring at Picomolar Concentrations with Automated Calibration

**DOI:** 10.1101/2025.06.13.659459

**Authors:** Claire M. S. Michielsen, Junhong Yan, Max H. Bergkamp, Yu-Ting Lin, Stijn R. R. Haenen, Rafiq M. Lubken, Alexander Gräwe, Anna Swietlikowska, Arthur M. de Jong, Menno W. J. Prins

## Abstract

Continuous monitoring of specific proteins is essential for understanding the dynamics of biological systems and for enabling real-time measurement-and-control strategies in bioprocesses. Ideally, sensors for continuous monitoring should be intrinsically reversible and able to perform accurate measurements over long time spans. Here, we present a particle nanoswitch sensor containing two different antibody fragments that bind reversibly to a protein of interest and thus form transient sandwich complexes. The antibody fragments are incorporated into the sensor using site-specific conjugation strategies to achieve optimal antibody orientation. Short-lived sandwich complexes are detected with single-molecule resolution, by tracking the motion of tethered particles. The sensing concept is demonstrated for lactoferrin, an iron-binding and immune-modulating protein. We show continuous measurements of picomolar concentrations with a response time of ∼10 min over periods of 12-15 h. Automated calibration strategies are described that result in a mean absolute relative difference below 10% compared to reference measurements. These results demonstrate how continuous fast protein sensing at picomolar concentrations can be achieved using reversible sandwich-based particle nanoswitches, enabling long-term monitoring of dynamic bioprocesses.

## Introduction

Proteins play key regulatory roles in biological systems and serve as essential markers for monitoring bioprocesses in fundamental biology, patient monitoring, and industrial bioprocessing^1-3^. Fluctuations of protein levels are conventionally determined through laboratory assays that analyze batches of samples collected at multiple time points. However, these laboratory-based methodologies have long turnaround times, typically a day or more, making it impossible to develop measurement-and-control strategies with rapid feedback to the bioprocess.

To enable bioprocess monitoring with rapid feedback, protein sensing technologies are needed which can operate continuously and have short response times. A promising approach in the field of continuous protein sensing is to design nanoswitches: nanotechnological constructs with a switching behavior that is modulated by the binding of a specific protein of interest^4–6^. Unlike traditional assays, nanoswitch-based sensors operate without consuming reagents, making them suitable for long-term use. An essential property of a nanoswitch for continuous monitoring is reversibility, in order to allow the monitoring of increases as well as decreases in protein concentration.

Recently, the reversibility of a molecular pendulum nanoswitch with picomolar sensitivity was achieved by applying an active reset mechanism^4^. The reset involved an applied voltage and current pulse that affects the protein interaction and regenerates the sensor by external intervention^4^. Alternatively, nanoswitch reversibility can be achieved by designing a nanoswitch based on molecular interactions that are intrinsically reversible^5,6^. Intrinsic reversibility has been achieved by incorporating binder molecules with sufficiently high dissociation rate constants, allowing bound proteins to rapidly and spontaneously unbind without any external intervention^5,6^. However, the sensitivity of these intrinsically reversible nanoswitches was limited to the nanomolar concentration range, underscoring the difficulty to achieve rapid intrinsic reversibility as well as high sensitivity.

In this study, we present an intrinsically reversible, sandwich-based particle nanoswitch sensor that achieves accurate protein quantification in the picomolar concentration range. The design incorporates antibody fragments (Fabs) that are conjugated to the particles and sensor surface using efficient and site-specific strategies. The Fabs have optimized kinetic properties that facilitate rapid and reversible protein sensing based on the detection of fast-dissociating sandwich complexes. The protein sensing principle is demonstrated for the monitoring of lactoferrin, as a model protein. Lactoferrin is an 80 kDa iron-binding glycoprotein that supports the immune system and is present in secretory fluids such as milk^7-9^. Continuous quantification of lactoferrin can enable real-time optimizations of processes to produce and purify such bioactive proteins. In this work, continuous protein sensing is demonstrated using automated calibration for accurate, real-time protein concentration measurements. The accuracy is assessed by studying signal variations and comparing sensor data to high-performance liquid chromatography (HPLC) measurements as a reference method. Finally, we discuss the potential of the nanoswitch-based sensing principle for real-time protein monitoring in bioprocessing applications.

## Results and Discussion

The molecular design and the readout methodology of the sandwich-based particle nanoswitch are illustrated in Figure 1A. The sensing cartridge is a microfluidic flow cell with a 7.5 µL chamber containing biofunctionalized particles, 1 µm in diameter, that are tethered to the substrate via a double-stranded DNA linker (dsDNA, 221 bp, contour length ∼75 nm)^10^. Both the particles and the polymer-coated substrate are functionalized with fast-dissociating Fabs. Several Fabs were screened for their ability to bind simultaneously to lactoferrin as a fast-dissociating sandwich pair (Supporting Information Figure S3). Based on the kinetic analysis from a previous study, Fab1 has a dissociation time on the order of minutes, and Fab2 has a dissociation time on the order of seconds (Fab1 and Fab2 correspond to Fab13 and Fab11 in reference^6^, respectively). A key function of the dsDNA tether is to keep the particle close to the surface, maximizing the encounter rate between Fab1 and Fab2. This ensures rapid sandwich formation in the presence of lactoferrin, leading to frequent transitions between bound and unbound states of the particle.

**Figure 1.**
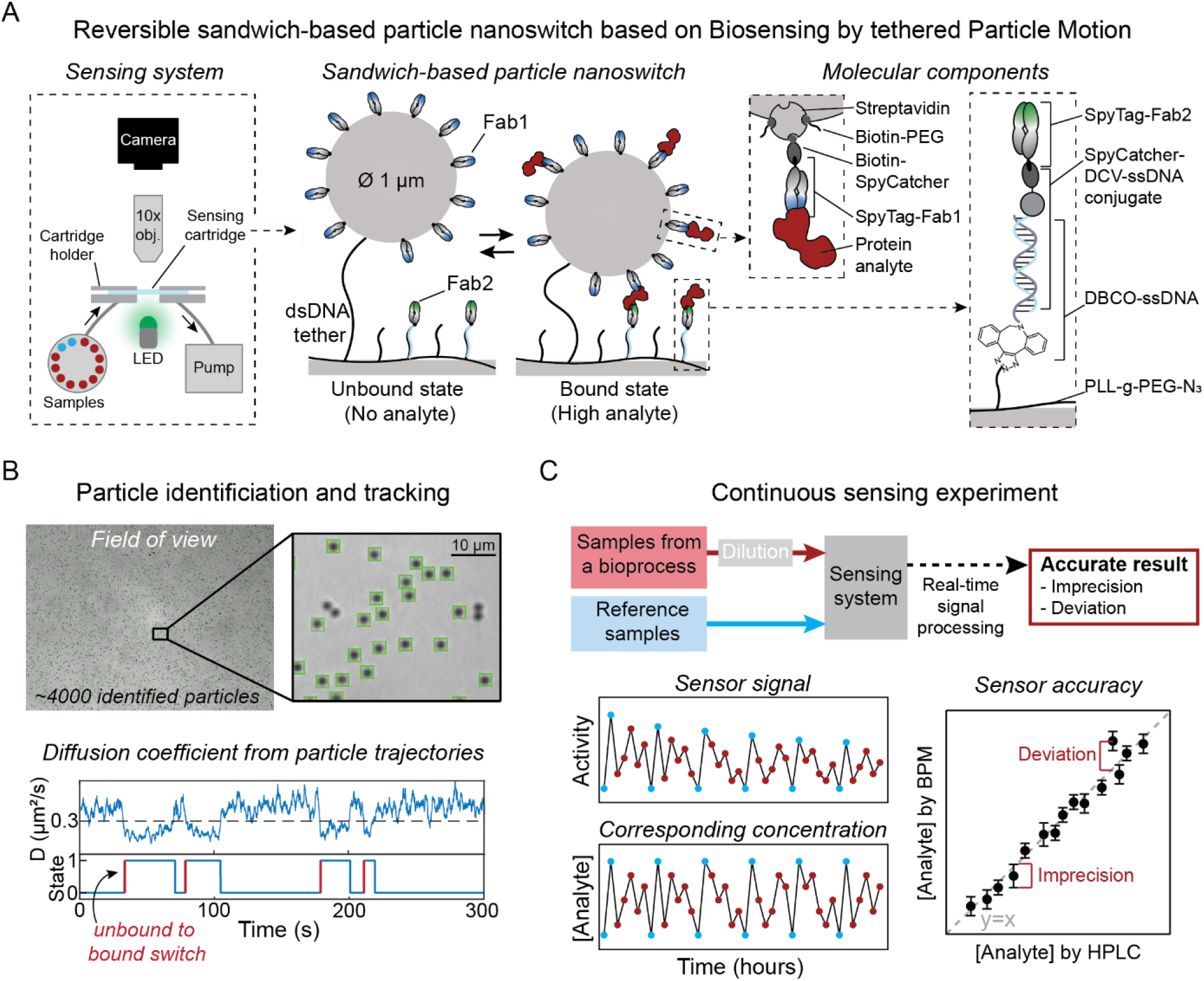
Design of reversible sandwich-based particle nanoswitch for continuous protein monitoring. (A) Schematic overview of the molecular design and the readout methodology of the particle nanoswitch. The sensing system comprises microfluidic components (tubing, valves, syringe pump), optical components (LED, objective, camera), a sensing cartridge and a computer for automated control of the fluidic and optical systems and for real-time signal processing. The sensing cartridge is a microfluidic flow cell with a 7.5 µL chamber containing biofunctionalized particles tethered to a biofunctionalized surface. In the presence of the protein analyte, the binders on the particles and the substrate form transient sandwich complexes with the analyte. Key molecular components are highlighted in the dotted-line boxes on the right. (B) Particle tracking and diffusivity-based signal processing. Green squares indicate particles that are identified by the tracking software, with approximately 4000 particles identified in the field of view. The software tracks the x and y positions of individual particles over time, generating particle trajectories. The diffusion coefficient is calculated from the particle trajectories and based on changes in the diffusion coefficient, transitions between bound and unbound states can be determined. The transitions from unbound to bound are used as the readout parameter. Details are given in Supporting Information Section S2. (C) Continuous protein sensing workflow. Diluted bioprocess samples (red) and reference samples (blue) are flown into the sensing system. The sensor signal is analyzed in real time to determine the corresponding protein concentrations. Sensor accuracy is assessed by evaluating concentration imprecision and by comparing sensor-based quantifications with HPLC results (deviation).

The total sensing system comprises the sensing cartridge with particles, fluidic components (tubing, valves, syringe pump), optical components (LED, objective, camera), and a computer for system control and real-time signal processing, see Figure 1A (leftmost panel). Video microscopy and particle tracking software are used to measure the motion of the particles, see Figure 1B. The software identifies individual particles and tracks their x- and y-positions over time, generating particle trajectories. From these trajectories, the diffusion coefficient is calculated as a function of time. The formation of an analyte-induced sandwich complex restricts the motion of the particle, leading to a reduced diffusion coefficient. The primary readout parameter is the frequency of switching between bound and unbound states. Previous publications on particle-motion based sensing (Biosensing by Particle Motion, BPM) used the total switching activity as the readout parameter, counting the frequency of all transitions: from bound-to-unbound as well as from unbound-to-bound^11–15^. In this work, only the transitions from unbound to bound are counted, referred to as Activity UTB (unbound to bound). Activity UTB provides a more robust measure of the sensor response than the switching activity based on all state transitions, as explained in Supporting Information Section S2. For the rest of the manuscript, the Activity parameter refers to Activity UTB.

The sensing workflow of this study involves the injection of a series of bovine milk samples from an industrial extraction process into the sensing system, along with reference samples of known lactoferrin concentration, into the sensing system (Figure 1C). The real-time signal processing software uses the reference samples for calibration to achieve accurate protein quantifications of blind samples. The sensor accuracy is assessed by evaluating concentration imprecision and deviations from the HPLC reference method.

### Molecular Components and Conjugation Strategies of the Nanoswitch

The molecular assembly of the nanoswitch is illustrated in Figure 2A. A cyclic olefin copolymer (COC) sensor surface was coated with a low-fouling polymer layer composed of PLL-g-PEG and PLL-g-PEG-azide. Single-stranded DNA (ssDNA) and dsDNA, both functionalized with DBCO, were conjugated to the azide groups on the PLL-g-PEG layer using click chemistry. Biofunctionalized particles, containing Fab1, were tethered to the surface via dsDNA, which contains a DBCO moiety on one end and a biotin moiety on the other end. To prevent multi-tethering, streptavidin-coated particles were partially blocked using biotin-PEG (1 kDa) after functionalization with Fab1. After particle tethering, an additional incubation step with biotin-PEG was used to block any remaining biotin binding sites.

**Figure 2.**
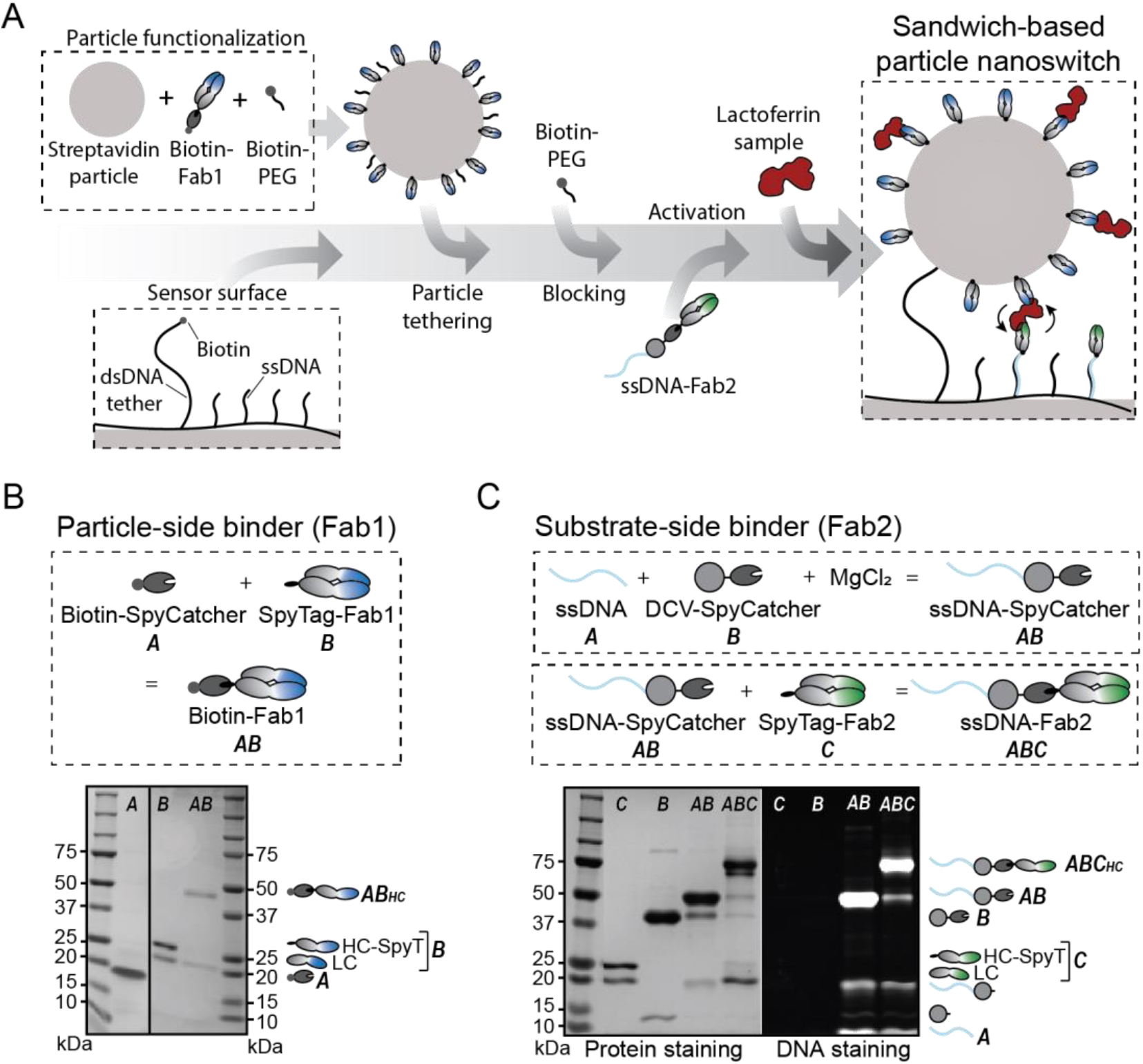
Molecular components and conjugation strategies of the nanoswitch. (A) Step-by-step assembly of the nanoswitch sensor. Particles are functionalized with biotin-Fab1 and partially blocked with biotin-PEG (1 kDa) to allow tethering. After attachment to the sensing surface via dsDNA tethers, the particles are further blocked with biotin-PEG. The sensing surface is coated with a low-fouling polymer containing both dsDNA tethers for particle attachment and ssDNA strands for secondary binder hybridization (ssDNA-Fab2). After adding the secondary binder, the nanoswitch sensor is ready for use. (B) Particle-side binder conjugation and SDS-PAGE analysis. Fab1, an antibody fragment containing a SpyTag, was conjugated to SpyCatcher-biotin, enabling site-specific biotinylation of Fab1. The SDS-PAGE gel displays the molecular components before (A and B) and after conjugation (AB). The Fab does not contain a disulfide bond between the heavy and the light chain, therefore two bands are observed^19^. (C) Substrate-side binder conjugation and SDS-PAGE analysis. The protein conjugate DCV-SpyCatcher was used to attach DNA to Fab2. Subsequently, Fab2, containing a SpyTag, was conjugated to the ssDNA-SpyCatcher complex, forming the ssDNA-Fab2 conjugate. SDS-PAGE analysis, with both protein and DNA staining, illustrates the molecular components before and after conjugation. The DCV-ssDNA reaction was incubated overnight at 4°C and the SpyCatcher conjugation was completed in 2 h at room temperature.

Fab1 was efficiently and site-specifically coupled to the particle using the SpyCatcher/SpyTag system^16^. SpyTag binds with high affinity to SpyCatcher, spontaneously forming a covalent isopeptide bond. A SpyTag fused to the heavy chain of Fab1 allows efficient conjugation of the antibody fragment to biotin-SpyCatcher, by mixing them together in PBS at a 1:1.3 ratio biotin-SpyCatcher:Fab-SpyTag and incubating for 2 h. This strategy enables controlled biotinylation without requiring a large reagent excess or subsequent purification steps. Figure 2B illustrates the molecular components, and SDS-PAGE analysis confirms successful conjugation.

Fab2, the secondary binder, was conjugated to the low-fouling polymer layer through DNA hybridization to ssDNA molecules, ensuring precise orientation and optimal binding efficiency. To attach ssDNA to Fab2, the protein conjugate DCV-SpyCatcher was used (Figure 2C). DCV, an endonuclease from Muscovy Duck Circovirus, selectively binds to a specific ssDNA sequence (TATTATTAC) in the presence of Mg^2+^, enabling efficient covalent and site-specific ssDNA-protein conjugation^17,18^. The conjugation reaction was performed using a 1:2 ratio of SpyCatcher-DCV to ssDNA. Subsequently, Fab2, which also contains a SpyTag, was conjugated to the ssDNA-SpyCatcher complex using a 1.3:1 ratio, resulting in the ssDNA-Fab2 conjugate. The DCV-ssDNA reaction was incubated overnight at 4°C, followed by the SpyCatcher-SpyTag reaction, which was completed within 2 h at room temperature. SDS-PAGE analysis with Coomassie staining was used to visualize proteins, and SYBR gold staining was used to visualize DNA, confirming the success of each conjugation step (Figure 2C). The majority of the DCV protein was conjugated to ssDNA, since only a small fraction of unconjugated DCV was observed. An impurity between 10–15 kDa was detected in the DCV-SpyCatcher sample, which shifted upward after DNA conjugation. This low-intensity band suggests that a small amount of DCV proteins lacking SpyCatcher was also expressed. We conclude from the data that the Fab2-SpyTag conjugation was efficient, as nearly all Fab2 reacted with SpyCatcher. This biomolecular assembly strategy allows covalent and site-specific conjugation without requiring a large excess of reagents.

### Continuous Protein Sensing Using Sandwich-Based Particle Nanoswitches

Continuous protein sensing was demonstrated using buffer samples spiked with picomolar concentrations of bovine lactoferrin (Figure 3). Each sample was flown into the flow cell at a flow rate of 100 µL/min for 1 min, after which signals were measured during 10 consecutive 1-min intervals in a static condition without flow. The blank measurement (sample without lactoferrin) yields a low background signal (Figure 3A). Increasing lactoferrin concentrations result in higher switching activities, due to the formation of protein-induced sandwich complexes. The measurements show that the sensor responds within a minute to changes in analyte concentration and that the signal was stable over the subsequent 10-min measurement period. After measuring the highest lactoferrin concentration (1000 pM), three blank samples were briefly flown into the flow cell, each followed by 10 min of measuring. The first and second blank samples bring the signals to the levels corresponding to 100 pM and 10 pM, respectively, with a signal relaxation time of a few minutes. The signal relaxation time is attributed to the slowest molecular dissociation process, i.e. the dissociation of lactoferrin from Fab1. The flow protocol (1-min flow duration) used in Figure 3 is estimated to replace approximately 90% of the fluid in the measurement chamber. In the following figures, two 1-min flow durations are used to ensure a carry- over of less than 10%.

**Figure 3.**
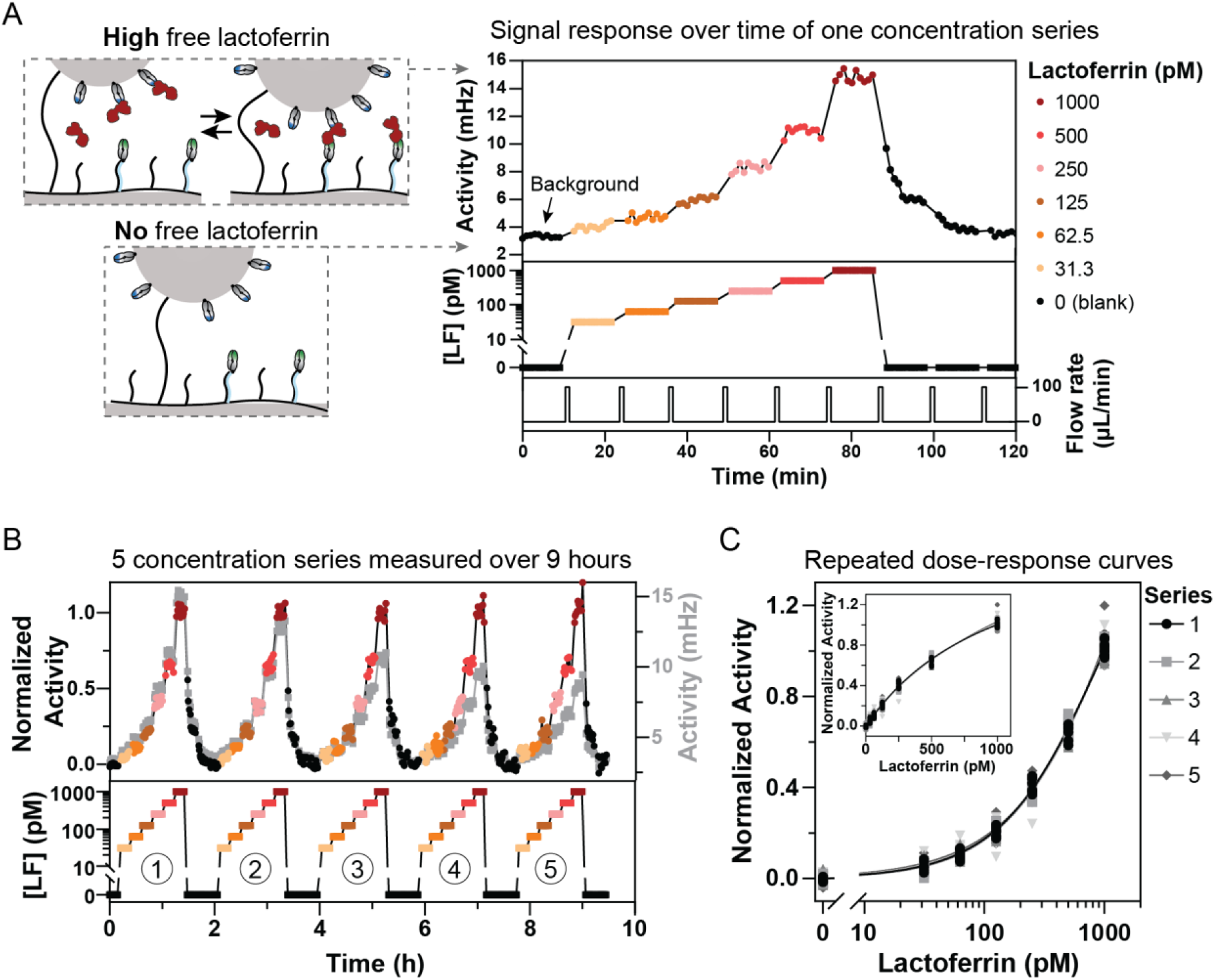
Continuous protein sensing using sandwich-based particle nanoswitches. (A) Sensor response over time for a single concentration series of lactoferrin. The switching activity of the sensor increases with higher lactoferrin concentrations and returns to background levels in the absence of lactoferrin, as illustrated in the schematics on the left. The top panel of the graph shows the sensor response and the middle panel the time profile of lactoferrin concentrations (0, 31.3, 62.5, 125, 250, 500, and 1000 pM in PBS with 500 mM NaCl and 0.1% BSA). The bottom panel illustrates the flow rate through the measurement chamber. Each sample was flown for 1 min at a flow rate of 100 µL/min, after which the measurement started immediately. The signal response was recorded under static conditions (no flow) using 10 consecutive 1-min intervals. (B) Continuous measurement of five series with six different lactoferrin concentrations over a 9-h period. Gray data points show the uncorrected activity signals (right y-axis). The colored and black data points show the normalized activity signals. (C) Dose-response curves of the five concentration series plotted with a logarithmic x-axis. The inset presents the data with a linear x-axis. Different symbols and shades of gray represent individual series, with all measured data points. Solid lines indicate the sigmoidal fits (Equation 1) per individual series, with fit parameters provided in Supporting Information Table S2.

To assess the measurement repeatability and sensor stability, the concentration series (31.3 pM to 1 nM) was measured five times over a 9-h period, see Figure 3B. The gray data points show the raw activity values (right y-axis), of which the signal gradually decreases over time. The decrease can be caused by mechanisms such as loss of binders on the particles, loss of binders on the substrate, and nonspecific interactions^20,21^. To correct for the gradual decrease of the signal, the data were normalized using linear interpolation between the highest and lowest signal points: the highest signal corresponds to the maximum analyte concentration (1 nM), and the lowest signal represents the background measurement (absence of analyte)^6^. The signal normalization leads to stable signal values across the serial measurements performed over 9 h, as shown in Figure 3B, and results in overlap of the five corresponding dose-response curves, see Figure 3C.

### Lactoferrin Concentration Measurements in Milk

The continuous nanoswitch sensor was further studied using milk as a biological fluid (Figure 4). Since the sensor is sensitive to picomolar concentrations, the milk samples were diluted 2000 times in order to bring the lactoferrin concentration into the sensitive range of the sensor. To evaluate sensor variability, six consecutive calibration cycles consisting of five different milk samples were measured on a single cartridge over a 13-h period (Figure 4A). Signal normalization was performed as described in the previous section: a milk sample with 125 pM lactoferrin was used to determine the zero signal of normalization, and a milk sample with 506 pM lactoferrin was used to determine the unity signal of normalization (the concentrations refer to the samples after dilution). The signal variations of all data points are shown in Figure 4B and the underlying distributions are shown in Supporting Information Figure S4. A calibration curve was fitted using all data points, as described by Equation 1:

**Figure 4.**
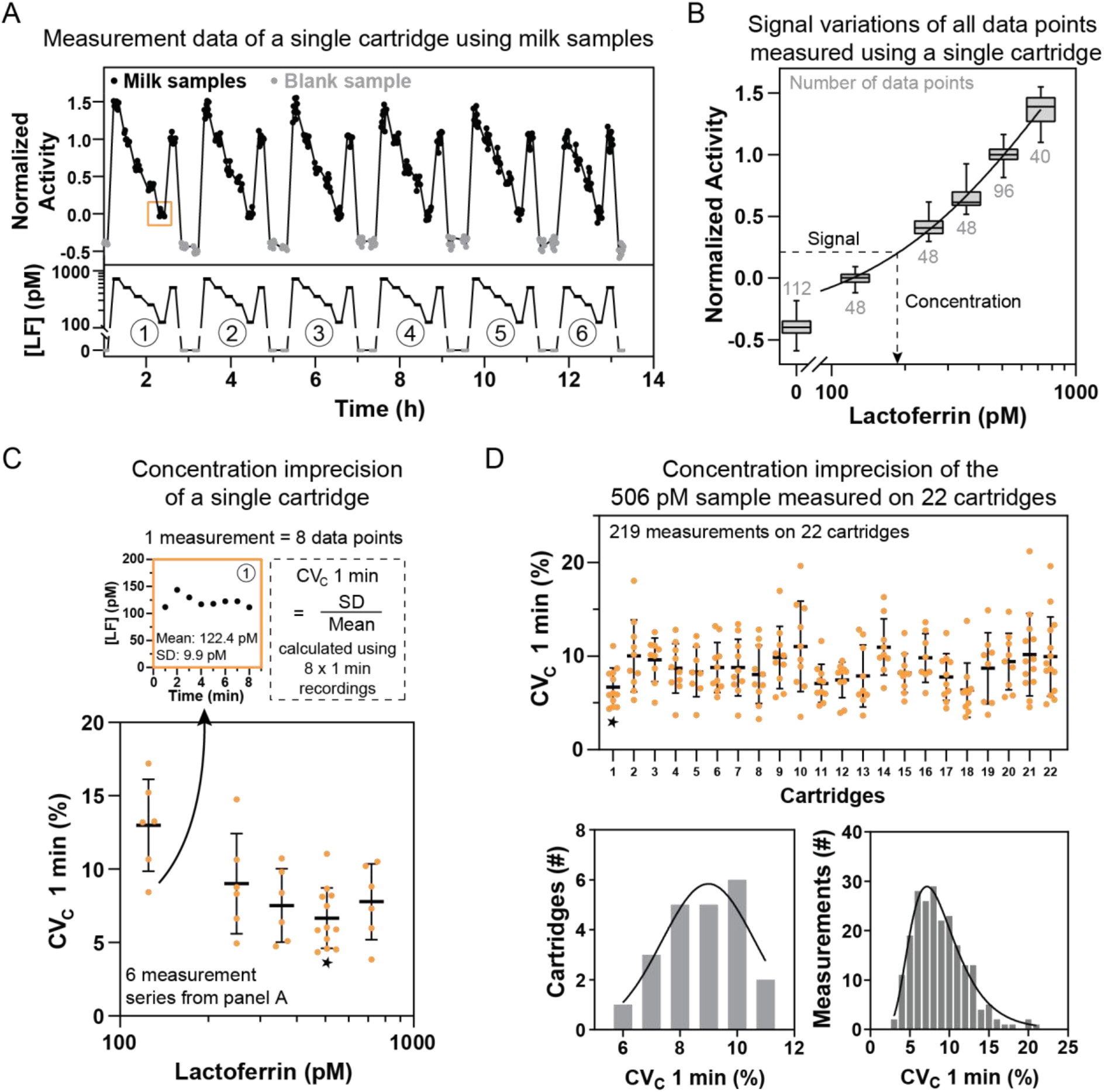
Concentration imprecision of lactoferrin measurements in milk, measured on multiple cartridges. (A) Measurement data from a single cartridge for five milk samples. The top panel of the graph shows measurement data (black data points) from five milk samples, one of which was measured twice per series (as indicated in the bottom panel). Milk samples were 2000x diluted in PBS with 500 mM NaCl. Blank samples (gray data points) were measured between different series. Six measurement series were conducted over approximately 13 h. Samples were flown into the cartridge using a flow rate of 100 µL/min for 1 min, the flow protocol was performed twice for efficient fluid exchange in the measurement chamber. (B) The number of data points per concentration is indicated in gray. Box plots illustrate the distribution of data points: whiskers represent the full range (minimum to maximum), the horizontal line within each box indicates the median and the box itself represents the interquartile range (50% of the data points). The solid line represents the sigmoidal fit of the dose-response curve, used to convert signal values into concentration values. (C) Concentration imprecision of a single cartridge. Each measurement consists of eight 1-min interval recordings, resulting in eight data points. The concentration imprecision, expressed as the coefficient of variation (CVC 1 min), was calculated by dividing the standard deviation by the mean of the eight data points. Orange data points represent individual measurements from panel A, the black horizontal line indicates the mean CVC 1 min per concentration of the repeated measurements on a single cartridge and the whiskers represent the standard deviation. (D) Concentration imprecision (CVC 1 min) of a single milk sample measured using multiple cartridges. A total of 219 measurements were performed on 22 different cartridges for a milk sample with a lactoferrin concentration of 506 pM. Orange data points represent the individual measurements, black horizontal lines represent the mean concentration imprecision of all measurements of the 506 pM sample on a single cartridge and the whiskers show the standard deviation. The histogram on the left displays the distribution of the mean concentration imprecision values per cartridge. The mean concentration imprecision per cartridge follows a normal distribution (p = 0.835, Anderson-Darling test) with a fitted mean and standard deviation of 9.0 ± 1.6%. The histogram on the right shows the distribution of the mean concentration imprecision values of all measurements. The concentration imprecision of all measurements follows a log-normal distribution (p = 0.342, Anderson-Darling test), with a geometric mean of 8.4% (5.7%-12.5%; 68 %-CI of the fit).

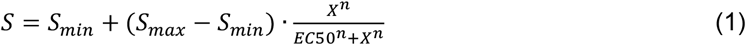

where *S*_*min*_ is the bottom plateau, *S*_*max*_ the top plateau, *X* the lactoferrin concentration, and *n* the slope factor. The calibration curve is used to convert signal values into concentration values.

The concentration imprecision was quantified as the coefficient of variation of 1-min recordings (CV_C_ 1 min), calculated by dividing the standard deviation of the eight data points per measurement (each representing a 1-min recording of the same sample) by the mean (Figure 4C). The average concentration imprecision ranged from 6.7 ± 2.0% to 13.0 ± 2.9% across the different concentrations. Imprecision was highest at the lowest analyte concentration, which can be attributed to the Poisson nature of the stochastic binding events. At low analyte concentrations, fewer binding events occur, leading to increased relative Poisson noise, and thus greater variability in the measured signals (Supporting Information Figure S5). Dilutional linearity was observed for dilution factors between 500 and 8000, indicating that the sensor is not sensitive to matrix differences within this range, see Supporting Information Figure S6. Therefore, reducing the dilution factor can be a strategy to minimize the imprecision of the samples with lower lactoferrin concentrations.

To study cartridge-to-cartridge variations, the concentration imprecision of one sample (506 pM) was evaluated across 22 cartridges. The sample was measured between 8 and 14 times per cartridge, each data point is shown in the top panel of Figure 4D. The histogram of the mean concentration imprecision per cartridge follows a normal distribution (p = 0.835, Anderson-Darling test), yielding a fitted normal distribution with a mean concentration imprecision of 9.0 ± 1.6%. The histogram of the concentration imprecision values of all measurements follows a log-normal distribution (p = 0.342, Anderson-Darling test), resulting in a geometric mean of 8.4% (5.7%-12.5%; 68%-CI of the fit).

When studying the variation over time for each cartridge, an increase in imprecision was observed. While the mean concentration remains constant across multiple measurements on the same cartridge, the variability increases over time. This can be explained by the gradual decrease in the dynamic range of the actual signal, which reduces the signal-to-noise ratio and causes larger fluctuations in the concentration quantifications, leading to greater imprecision in later measurements. The analysis of concentration imprecision over time for different cartridges is provided in Supporting Information Figure S7.

### Calibration Strategies for Accurate Quantification

To obtain accurate concentration quantifications of blind milk samples, different calibration strategies were evaluated. Figure 5A presents the measurement data of reference and blind milk samples. The reference samples are milk samples with a known concentration of lactoferrin determined using HPLC. A full calibration curve was generated using five reference samples, two of which were measured approximately every two hours for signal normalization. Four blind milk samples were measured in varying orders, with the two normalization reference samples included after every five blind sample measurements. Blank samples were used as additional controls to monitor the sensor background over time. Based on this dataset, different quantification strategies using Equation 1 were evaluated (Figure 5B): (1) performing a repeated calibration fit using five reference concentrations, with recalibration every four hours; (2) conducting a single calibration fit with five reference concentrations at the beginning and using this calibration fit for all subsequent measurements; or (3) calibrating via a fixed calibration curve, which was determined based on the data from Figure 4B (Equation 1 with S_min_ = −0.5, S_max_ = 4, EC50 = 1.03 nM and n = 1), to quantify the blind samples. For all three methods, the signals were normalized using the normalization reference samples.

**Figure 5.**
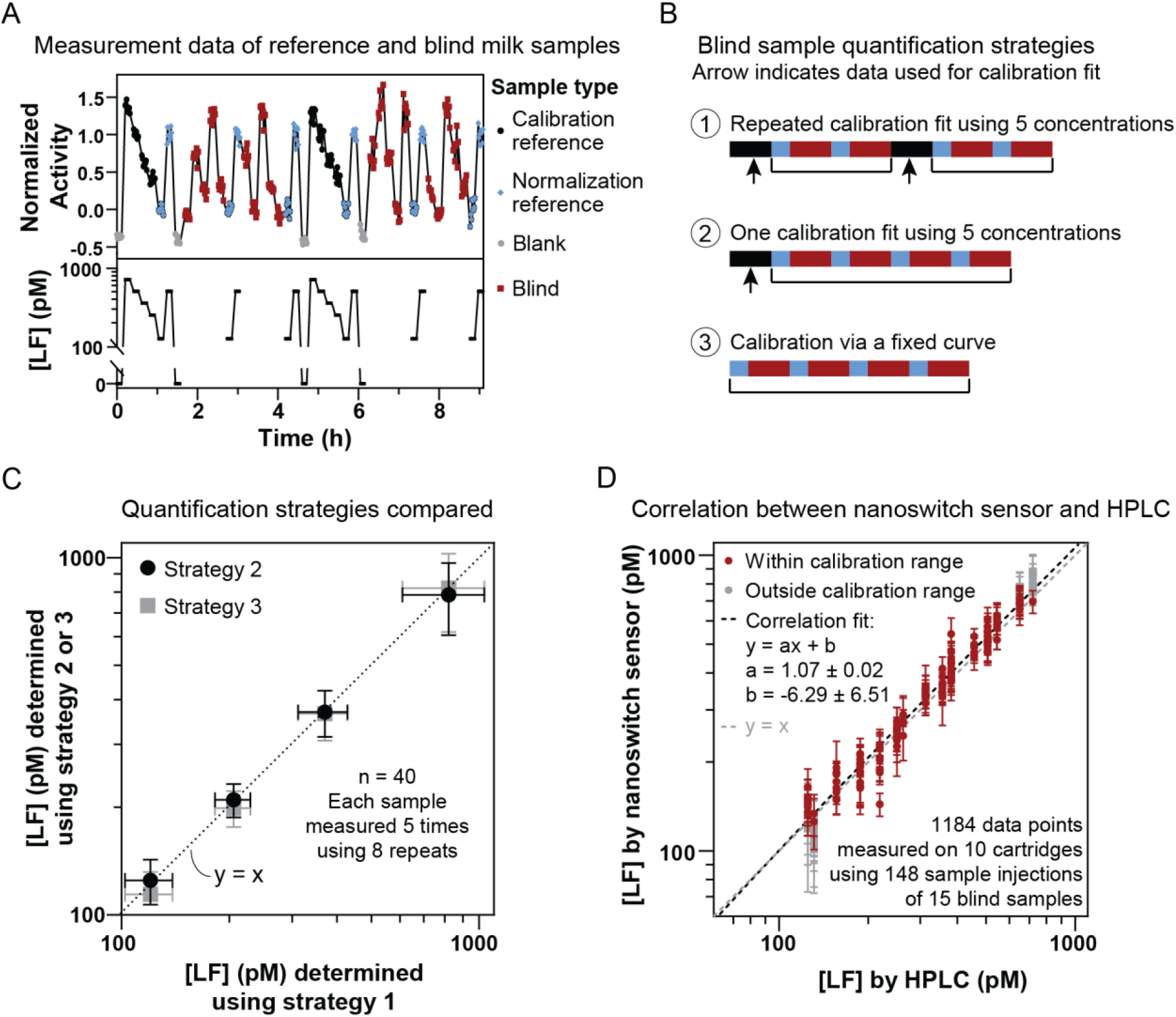
Automated calibration for accurate protein quantification in milk. (A) The top panel of the graph shows the measurement data and the bottom panel displays the lactoferrin concentrations of the reference samples. Reference milk samples used for calibration are shown in black, reference milk samples for data normalization in blue, blank buffer samples in gray, and blind milk samples in red. Four different blind samples were measured repeatedly in varying orders. All milk samples were 2000x diluted in PBS with 500 mM NaCl. (B) Three strategies for lactoferrin quantification were used. Strategy 1 involved repeated calibration fits using five reference concentrations. Strategy 2 applied a single calibration fit based on five reference concentrations measured at the beginning of the experiment. Strategy 3 used a fixed calibration curve. For all three methods, the data was normalized using the normalization reference samples, indicated in blue. (C) Comparison of the three quantification strategies using data from panel A. Concentrations determined using strategy 1 are plotted on the x-axis, while concentrations determined using strategy 2 or 3 are plotted on the y-axis. Strategy 2 results are represented by black circles and strategy 3 results by gray squares. Each circle or square represents the mean of 40 data points per sample, with error bars indicating the standard deviation. (D) Correlation between lactoferrin concentrations determined by the nanoswitch sensor and by HPLC. Measurements within the calibration range are shown in red, while measurements outside this range are shown in gray and excluded from the linear correlation fit (black dotted line). The data points represent the mean concentration from eight 1-min recordings, with error bars indicating the standard deviation. The gray dotted line represents the Y = X correlation.

Quantification strategies 2 and 3 are evaluated against strategy 1 in Figure 5C. All three strategies yield similar results, indicating that calibration time can be saved by using the most efficient strategy number 3. Supporting Information Figure S8 gives further information about the three strategies using data from 10 cartridges, showing that a fixed calibration curve could be consistently applied across all cartridges, demonstrating high reproducibility. The ability to use a single calibration curve across multiple cartridges highlights the robustness and reproducibility of the sensing principle.

To evaluate the level of agreement between the nanoswitch sensor and HPLC measurements, the correlation between the two methods was analyzed, see Figure 5D. The results showed a linear relationship between the two methods, with the dotted gray line representing the Y = X correlation and the dotted black line indicating the linear fit of the data. Data points outside the calibration range were excluded from the fit. The two correlation lines closely align, confirming the linearity of the correlation and the agreement between the two methods. The relative deviations from the Y = X line are shown in Supporting Information Figure S9. To summarize the analytical performance of the sensor in a single value, we calculate the mean absolute relative difference (MARD), which quantifies the agreement between the nanoswitch sensor and the HPLC measurements. The MARD is 8.2% for the concentration range from 125 to 719 pM (n = 122). Both the imprecision, quantified as CV_C_ 1 min, and the deviation from reference measurements, quantified as MARD, are below 10% for the proof-of-concept continuous sandwich-based protein sensor studied in this paper.

## Conclusion

Intrinsically reversible nanoswitches are a promising approach to enable the continuous sensing of biomolecules. In this study, we have developed a sandwich-based particle nanoswitch for continuous protein monitoring with single-molecule resolution. The use of site-specific conjugations of fast-dissociating antibody fragments generated via phage display methodologies^19,22^, makes the approach versatile for a wide range of applications. The nanoswitch sensor was evaluated for monitoring lactoferrin in milk using an automated calibration methodology. The sensor achieves picomolar sensitivity with a response time of approximately 10 min and shows stable operation over 12-15 h.

A key innovation in the developed reversible nanoswitch sensor is the detection of transient sandwich complexes with single-molecule resolution. Antibody fragments Fab1 and Fab2 were selected for their ability to bind simultaneously to the target protein as a short-lived sandwich pair. The dsDNA tether between particle and surface induces a high encounter rate between the Fab1 and Fab2 molecules, which ensures rapid sandwich formation in the presence of lactoferrin and causes frequent transitions between bound and unbound states of the particle.

The sandwich-based nanoswitch sensor continuously measures protein concentrations with a fast response time, without needing any regeneration. This is caused by the fact that protein molecules continuously bind and unbind at high rates. The nanoswitches reach a dynamic equilibrium, with a relatively low fractional occupation of Fabs by proteins, due to high dissociation rates and low protein concentrations. The dynamic equilibrium involves continuous stochastic binding and unbinding of proteins, resulting in continuous stochastic switching of the particles between bound and unbound states, due to the transient formation of single sandwich complexes between particle and surface. The well-defined molecular interactions lead to dose-response curves with consistent shapes, allowing for automated calibrations with only two concentration references.

The imprecision, quantified as CV_C_ 1 min (coefficient of variation of protein concentration determinations from 1-min recordings), and the deviation from reference measurements, quantified as MARD (mean absolute relative difference of concentration determinations), are both below 10% for the nanoswitch sensor studied in this paper. A fundamental contributor to the variability of the nanoswitch sensor signals relates to the stochastics and counting statistics of the particle switching events, originating from single-molecular binding and unbinding events. We hypothesize that the counting statistics are the dominant factor in the observed precision (see Supporting Information Section S6). Further work will focus on disentangling the origins of the variations and studying how these scale with statistics-related parameters such as the number of particles, the observation time, association and dissociation rate constants, and densities of Fabs on particle and substrate.

The demonstrated accuracy of the nanoswitch-based protein sensor can be compared to the concentration fluctuations in bioprocessing applications. In this paper, lactoferrin was used as a model protein target. Lactoferrin is a bio-active protein that is produced by extraction and purification from bovine milk^23,24^ and can also be produced by precision fermentation^25^. In bovine milk, lactoferrin concentrations vary by several tens of percents due to multiple independent factors, including stage of lactation, milk yield and somatic cell count^26–29^. Such fluctuations propagate into the extraction and purification processes of the protein. In precision fermentations, the concentration fluctuations are even larger, in upstream as well as in downstream processing. Therefore, the demonstrated analytical performance of the proof-of-concept sensor is already at a useful level for bioprocessing applications.

To our knowledge, the presented nanoswitch sensor is the first continuous protein sensor that is intrinsically reversible and can measure in the picomolar range. The nanoswitch sensor does not consume any reagents, which makes the sensing method cost-effective and suited for long-term use. Future research will explore the sensitivity limits, study binder designs and nanoswitch engineering using computational methods^30–34^, enhance sensor stability for operation over days and weeks, and expand the sensor’s capabilities for continuous sensing of a wide range of protein markers that can impact the fields of bioprocessing, healthcare, and environmental monitoring.

## Acknowledgements

We thank Royal FrieslandCampina (RFC) for providing the milk samples and HPLC measurement data. We thank Allard Smedinga and Hendrik Kleefstra for performing measurements at RFC. Part of this work was funded by The Netherlands Topsector Agri&Food, HTSM, and Chemistry under contract number LWV20.117. Part of this work was funded by The Netherlands National Growth Fund Programme NXTGEN HighTech.

