## Supporting Information for "Reversible Sandwich-Based Particle Nanoswitch for Continuous Protein Monitoring at Picomolar Concentrations with Automated Calibration"

### Table of Contents

S1. Material and Methods ... 2
S2. Biosensing by Particle Motion readout parameters ... 4
S3. Fast-dissociating sandwich pair screening in f-BPM ... 7
S4. Fitting parameters of dose-response curves in Figure 3C ... 8
S5. Signal variations for different lactoferrin concentrations in milk samples ... 8
S6. Influence of Poisson noise on concentration imprecision ... 9
S7. Dilutional linearity ... 10
S8. Concentration imprecision over time ... 11
S9. Calibration strategies compared ... 11
S10. Relative difference between nanoswitch and HPLC measurements ... 12
References ... 13

#### S1. Material and Methods

***Materials.*** Bovine lactoferrin powder and milk samples containing lactoferrin were supplied by FrieslandCampina. The custom HuCAL recombinant antibodies (Fabs) against bovine lactoferrin were produced by Bio-Rad[1,2]. The oligonucleotides used in this study are listed in Table S1 and were purchased from IDT. Custom-made cyclic olefin copolymer (COC) cartridges containing a microfluidic flow cell, compatible with the custom-built automated setup, were produced by Axxicon using injection molding. The flow cell has a height of 250 µm and a surface area of 80.5 mm$^2$, resulting in a total volume of 20.1 µL. The chamber containing the particles has a volume of 7.5 µL.

*Table S1. DNA sequences used in the nanoswitch sensor.*

| Name | Sequence |
|---|---|
| **dsDNA tether** | 5′ DBCO-GGT TAG CAG CCT GTT TCA AAA CCT GGG GGT GAG TGT CAC GCC AAT TCA GCG CAT CGT TCT GTC GGG AGA GAA TGG TCT GAA AAT CGA TAT CCA CGT CAT TAT CCC GTA CGA AGG TCT TTC TGG TGA TCA GAT GGG GCA GAT AGA AAA AAT ATT CAA AGT GGT GTA CCC AGT AGA CGA TCA TCA CTT CAA GGT TAT ACT GCA CTA TGG CAC CCT CGT TAT CG 3′<br><br>3′ CCA ATC GTC GGA CAA AGT TTT GGA CCC CCA CTC ACA GTG CGG TTA AGT CGC GTA GCA AGA CAG CCC TCT CTT ACC AGA CTT TTA GCT ATA GGT GCA GTA ATA GGG CAT GCT TCC AGA AAG ACC ACT AGT CTA CCC CGT CTA TCT TTT TTA TAA GTT TCA CCA CAT GGG TCA TCT GCT AGT AGT GAA GTT CCA ATA TGA CGT GAT ACC GTG GGA GCA ATA GC-biotin 5′ |
| **DBCO-ssDNA** (docking DNA) | 5′ DBCO-GTGCGGCAGGGGTAAGACCA 3’ |
| **DCV-ssDNA** (complementary to docking DNA) | 5' TATTATTACTTTTTTGGTCTTACCCCTGCCGCAC 3' |

***SpyCatcher003-DCV expression.*** SpyCatcher003-DCV was cloned via restriction and ligation into pET28a from existing plasmids in the Merkx group at Eindhoven University of Technology. For protein expression, E. coli BL21 (DE3) was transformed with the plasmid and plated on LB agar supplemented with 50 µg/mL kanamycin. Next day, a small overnight culture was inoculated with a single bacterial colony, which was inoculated into a large (750 mL) culture of LB medium supplemented with kanamycin the day after. The culture was incubated at 37 °C with vigorous shaking and induced with 0.5 mM IPTG upon reaching an OD600 of 0.6-0.8. Protein expression was performed overnight at 20 °C. Cells were harvested by centrifugation and lysed with BugBuster (Novagen) supplemented with benzonase (Merck) and cOmplete protease inhibitor tablet (Roche), and left shaking for 1 h at ambient temperature (21 °C). After centrifugation to remove cell debris, the supernatant was loaded on a StrepTactin-XT gravity flow column equilibrated with buffer W (150 mM NaCl, 100 mM Tris-Cl pH 8, 1 mM EDTA). To remove excess DNA, the column was then washed with high salt buffer (500 mM NaCl, 20 mM Tris-Cl pH 8, 10 mM Imidazole) with 10 column volume (CV), followed by a final 10 CV wash with buffer W without EDTA. The protein was eluted with buffer E (150 mM NaCl, 100 mM Tris-Cl pH 8, 50 mM D-biotin), checked for purity via SDS-PAGE, and aliquots were flash frozen in liquid N2 and stored at -80 °C.

***Endonuclease (DCV) ssDNA conjugation.*** SpyCatcher003-DCV was conjugated to ssDNA (sequence provided in Table S1) at a 1:2 molar ratio in the conjugation buffer (50 mM HEPES, pH 7.5, 50 mM NaCl, and 1 mM $MgCl_2$). The reaction mixture was incubated overnight at 4 ºC. Next day, the Fab-SpyTag/SpyCatcher conjugation was performed as described below. The third generation of this system (SpyTag003/SpyCatcher003) has been reported to exhibit near-diffusion limited kinetics.

***Fab-SpyTag/SpyCatcher conjugation.*** The SpyTag/SpyCatcher technology was used for site-directed conjugation of the recombinant Fabs containing a SpyTag002[1]. Fab1 (particle-side binder) was conjugated to Biotin-SpyCatcher002 (TZC001B; Bio-Rad) and Fab2 (substrate-side binder) was conjugated to the SpyCatcher003-DCV-ssDNA conjugate (described above). The conjugation reaction was performed in PBS at RT for 2 h, using a Fab-SpyTag002 to Biotin-SpyCatcher001 or SpyCatcher003-DCV-ssDNA molar ratio of 1.3:1. The conjugated Fabs were aliquoted and stored at -20 ºC.

***SDS-PAGE analysis.*** Protein samples were prepared by mixing 15 µL of sample with 5 µL of 4× Laemmli loading buffer (Bio-Rad) and incubating at 95°C for 10 min. SDS-PAGE was performed using Novex WedgeWell™ 4–20% Tris-glycine gels (Thermo Fisher) and Tris/glycine/SDS running buffer (Bio-

Rad). The protein samples and Precision Plus Protein™ All Blue standard (Bio-Rad) were loaded into the wells, and electrophoresis was carried out at 150 V for 1.5 h. After electrophoresis, the gel was first washed with Milli-Q water for 30–60 min. If both DNA and protein staining were performed, DNA staining was carried out first. The gel was incubated with 5 µL of 1000× SYBR™ Gold stain (Thermo Fisher) for 30 min at RT, followed by a 30-min wash with Milli-Q water. The gel was then imaged using the Bio-Rad Gel Doc EZ Imager. For protein staining, the gel was incubated with Coomassie Blue staining buffer (Bio-Rad) for 20–30 min at RT, followed by an overnight wash with Milli-Q water. The gel was visualized using the same imager.

**Continuous sandwich-based particle nanoswitch sensor**

***Particle Functionalization.*** 4 µL streptavidin-coated Dynabeads MyOne C1 (10 mg/mL; Thermo Fisher) were incubated with 8 µL 125 nM biotin-Fab1 for 30 min at RT on a rotating fin. The remaining biotin binding sites were partly blocked by adding 2.5 µL 10 µM biotin-mPEG (PG1-BN-5k, Nanocs) and 10 µL PBS. The mixture was incubated for 45 min at RT on the rotating fin, followed by washing with 1 mL PBST. Particles were either resuspended in HS buffer with 0.1% BSA for few-day storage, or were resuspended in sugar solution (25% sucrose and 100 mM trehalose in MilliQ water) and then dried for long-term storage.

***Cartridge Preparation.*** A polymer mixture of 0.45 mg/mL poly(L-lysine)-grafted-poly(ethylene glycol) (PLL(20)-g[3.5]-PEG(2); SuSoS) and 0.05 mg/mL azide functionalized PLL-g-PEG (PLL(15)-g[5]-PEG(2)-N3; Nanosoft Biotechnology LCC) was prepared in ultra-pure MQ, as described by Lin et al[3]. The COC cartridges were rinsed twice with Milli-Q water and cleaned using 15 min of sonication in Milli-Q water. The cartridges were dried using a nitrogen steam and exposed to UV Ozone (UV Ozone Cleaner, Novascan) for 30 min. Immediately after the UV Ozone treatment, the flow chamber of the cartridge was sealed with an adhesive film (391-0189, VWR) and the PLL-g-PEG/PLL-g-PEG-N3 mixture was injected. After 3 h of incubation at RT in a humidity chamber, the free polymer was removed by extracting the solution from the chamber. Subsequently, a DNA mixture was added, containing 0.4 nM dsDNA tether (DBCO modified) and 3 µM DBCO-ssDNA (docking DNA) diluted in HS buffer (PBS with 500 mM NaCl). The inlet and outlets of the flow chamber were sealed, and the cartridges were stored for at least 3 weeks, up to 3 months.

***Cartridge Activation.*** Particles in HS buffer with 0.1% BSA, or dried particles resuspended in HS buffer, were incubated in the cartridge for 15 min, to allow binding of the particles to the biotin moiety on the substrate-side dsDNA tether. From this point on, all fluid additions were done automatically at a flow rate of 100 µL/min and a flushing volume of 100 µL. After particle tethering, the remaining biotin-binding sites were blocked with 200 µM 1 kDa biotin-mPEG for 10 min. The system was activated using 25 nM Fab29-ssDNA, which hybridizes to the substrate-side docking DNA. After 20 min incubation of Fab29-ssDNA, the activation process was stopped by flushing HS buffer. At that point the cartridge was ready to measure samples with lactoferrin.

***Milk sample preparation.*** Milk samples were thawed at room temperature and briefly vortexed. Samples were diluted 2000 times in HS buffer, aliquoted in 4 mL glass vials, and stored in the freezer for up to a few months. Before each measurement, the frozen samples were thawed at 37˚C in a thermal shaker for 15 min. Samples were homogenized by inverting the vial a few times and then added to the measurement setup.

***Automated setup and Data Analysis.*** A Laboratory Programmable Syringe Pump (LSPone) and 12-port rotary valve from AMF were used to transport fluids. PTFE tubing (BL-PTFE-1608-20) and connectors (CIL-XP-245X) from Darwin Microfluidics were used to connect the LSPone pump and valve to the cartridge. A cartridge holder was custom-made from aluminum. O-rings were used for establishing watertight connections between holes in the cartridge holder and inlet and outlets of the cartridge. The particles inside the flow chamber were tracked using a custom-made optical setup containing a 10 times magnification objective (10x DIN achromatic finite intl standard objective; Edmund Optics), a 3 mm green led (12V), and a 3.2 MP camera (Flir BFS-U3-32S4M-C) with a field of view of 0.71 mm x 0.53 mm. The camera pixel size is 3.5 µm, which corresponds to a distance of 345 nm in the field of view. A miniature linear actuator (Zaber T-LA13A) was used for the autofocus. The complete setup (pumps, valves and microscope) was controlled by a computer. Measurements of 1 min at a framerate of 30 Hz were performed after a sample was flown through the flow chamber, so in the absence of flow. The frames of each measurement were analyzed in real-time using particle tracking software described by Bergkamp et al[4].

#### S2. Biosensing by Particle Motion readout parameters

Biosensing by Particle Motion (BPM) relies on reversible interactions between biofunctionalized particles and a biofunctionalized sensing surface (Figure S1A). The measurements are based on tracking the motion of the particles. When the particles are not interacting with the surface, they move freely, exhibiting high diffusion coefficients. In contrast, interactions with the surface limit the particle motion, resulting in lower diffusion coefficients. Using brightfield video microscopy and particle tracking software, hundreds to thousands of particles are analyzed simultaneously, allowing for robust statistical measurements[4].

To quantify the response of a BPM sensor with tethered particles (t-BPM), different readout parameters can be used. The two primary parameters are activity and bound fraction. Bound fraction is determined from the diffusion coefficient histogram of all particles in a time interval (1 min in this article) and represents the fraction of time particles spend in the bound state in the time interval (Figure S1B). Activity, on the other hand, quantifies the average number of state transitions per tracked particle per unit of time (Figure S1C). Previous BPM studies have used total switching activity as the readout parameter, which counts all transitions between bound and unbound states[5–8]. However, in this work, only transitions from the unbound to the bound state are considered. This refined metric is referred to as Activity UTB (Unbound to Bound). Activity UTB provides a more robust measure of the sensor response compared to total switching activity. By performing state classification and using only transitions from unbound to bound, Activity UTB reduces background and minimizes contributions from multivalent and nonspecific interactions.

The suitability of Activity UTB as a readout parameter depends on the interaction kinetics within the sensor. In a previously studied competition-based t-BPM sensor[9], the interactions were too slow to reliably use Activity UTB. In that case, bound fraction was preferred as the readout parameter. In the present study, the particles exhibit short-lived binding interactions. This increases the number of binding and unbinding events, providing sufficient statistics to use Activity UTB as the readout parameter.

Figure S2 shows the behavior of different readout parameters for the data presented in Figure 3. The bound fraction parameter shows an increase in the background and a decrease in the maximum signal as a function of time. The increase in the background is attributed to nonspecific interactions or multivalent interactions that cause particles to stay in the bound state. The decrease in the maximum signal is attributed to a loss of binders on the particles, loss of binders on the substrate, and nonspecific interactions[10,11].

The activity parameter shows a decrease in the background and a decrease in the maximum signal. The decreasing background can be attributed to particles that become stuck and stop switching. The decrease in the maximum signal is attributed to a loss of binders on the particles, loss of binders on the substrate, and nonspecific interactions[10,11].

The Activity UTB parameter shows a stable background signal and a decrease of the maximum signal. Compared to the activity, the background is more stable and lower, which is caused by the fact that the parameter counts only unbound-to-bound transitions, so it is not affected by particles that become stuck to the surface due to multivalent or nonspecific interactions. This makes Activity UTB a more robust readout parameter compared to bound fraction and total switching activity.

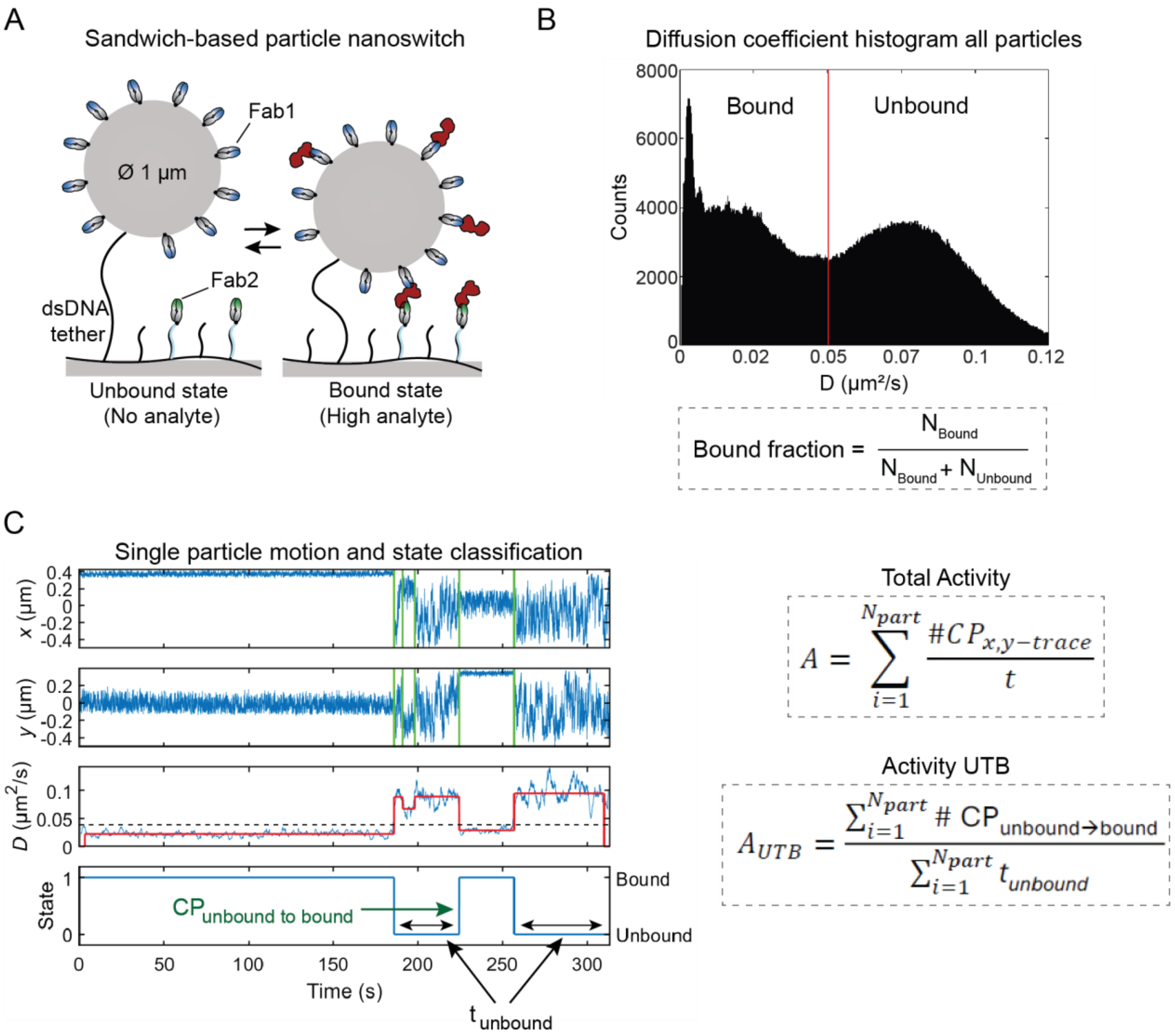


***Figure S1. Biosensing by tethered Particle Motion (t-BPM) sensing principle.*** *(A) Schematic of a single particle in a sandwich-based t-BPM sensor. The particle and substrate are coated with molecular binders, which form short-lived sandwich complexes in the presence of the target protein. The dsDNA tether keeps the particles in close proximity to the substrate, enhancing the association rate of the sandwich complexes. (B) Diffusion coefficient ($D$) histogram of all particles in the t-BPM sensor in a time interval of 1 min. In the presence of the analyte, the motion of the particle is restricted due to sandwich complex formation between the binders on the particle and substrate, resulting in low diffusion coefficients ($D < 0.05\ \mu m^2/s$). In the absence of the analyte, particles are in the unbound state, resulting in higher diffusion coefficients ($D > 0.05\ \mu m^2/s$). The bound fraction represents the fraction of time particles spend in bound states in the time interval. The red line represents the threshold used to determine the bound fraction. (C) Particle motion and state classification of a single particle. The top two panels show the x and y positions of the particle over time. Green vertical lines indicate change points (CP), which are used to calculate the total activity, the average number of CPs per particle per unit of time. Based on the x and y positions, the diffusion coefficient is calculated over time. The dotted line represents the diffusion coefficient threshold used for the classification of bound and unbound states. Allowing the identification of change points from unbound to bound, indicated with the green arrow. Activity UTB considers only switches from unbound to bound and the time spend in unbound states instead of the total measurement time.*

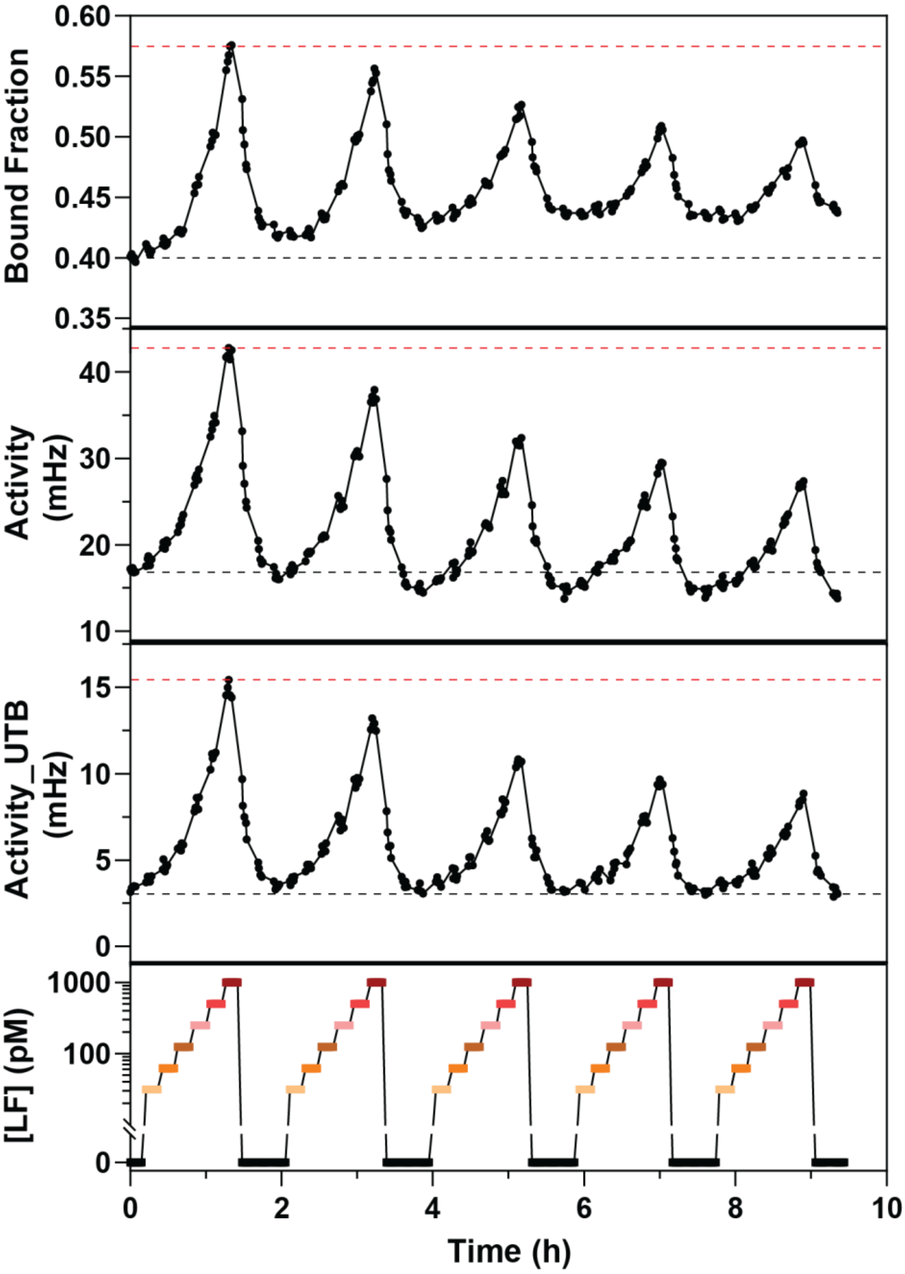


***Figure S2. Bound fraction, total switching activity, and activity_UTB readout parameters, determined from the data in Figure 3.*** *The top panel shows the bound fraction over time, the second panel shows the total switching activity over time, and the third panel shows Activity UTB over time. The bottom panel displays the lactoferrin concentrations applied to the sensor over time.*

#### S3. Fast-dissociating sandwich pair screening in f-BPM

Thirteen recombinant antibody fragments (Fabs) were characterized using Biosensing by Particle Motion (BPM) and Surface Plasmon Resonance (SPR) in a previous study[9]. Fast-dissociating Fabs were identified and were further screened to find a suitable sandwich pair. The screening was performed using free BPM (f-BPM) in a 96-well plate format. In f-BPM, particles are not tethered to the surface, which simplifies sample preparation and makes the method suitable for screenings in 96-well plates (Figure S3A). For f-BPM, the bound fraction was used as the readout parameter.

In the screening setup, streptavidin-coated particles were functionalized with various Fabs and the surfaces of the wells were coated with neutravidin and functionalized with Fab 11 (Figure S3A). Fab 11 was chosen as it exhibited the fastest dissociation in the SPR analysis[9]. The results are shown in Figure S3B. In the absence of lactoferrin, Fab 6 exhibited a very high background, and Fabs 5 and 7 also showed some background, making them less suitable for a sandwich sensor. Fab 11 on the particle (the same Fab as on the substrate), showed minimal sandwich binding, with only a slight background observed.

Fabs 8, 12, and 13 exhibited low background in the absence of lactoferrin and showed an increase in bound fraction as the lactoferrin concentration increased. Among these, Fab 12 and Fab 13 demonstrated better sensitivity compared to Fab 8. Fab 12 and Fab 13 were therefore identified as the most suitable candidates for a sandwich pair with Fab 11.

For the sandwich sensor in this study, Fab 13 was used as Fab 1 on the particle and Fab 11 as Fab 2 on the substrate.

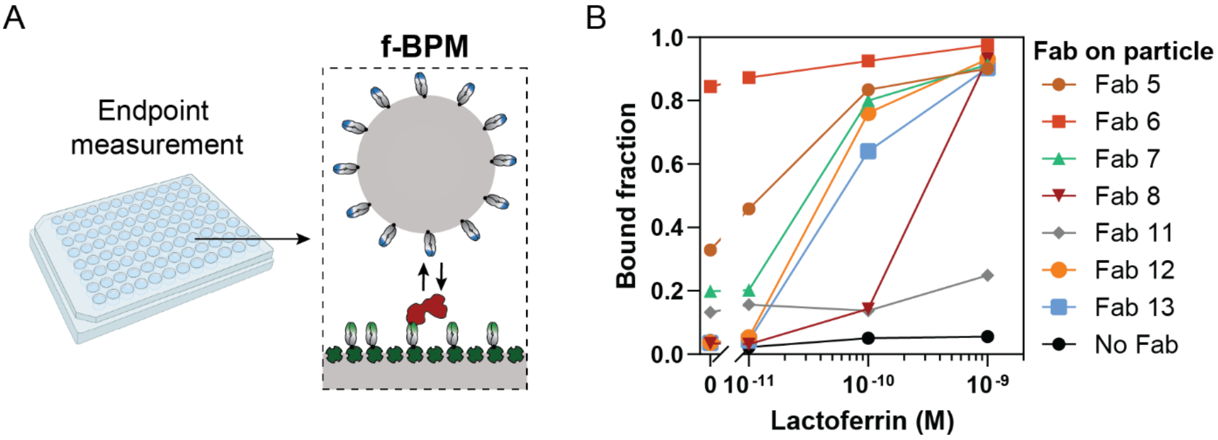


*Figure S3. Sandwich pair screening using Biosensing by free Particle Motion (f-BPM). (A) Schematic representation of the f-BPM sensor in a 96-well plate. The surface was functionalized with neutravidin and biotin-Fab 11, and different biotin-Fabs were coupled to streptavidin-coated particles (1 µm diameter). In the presence of the analyte lactoferrin, particles can form sandwich complexes that restrict the motion of the particles. (B) Dose-response curves for 7 Fabs, each represented by a different color and symbol. The Fab numbers correspond to the Fabs in the previous study[9]. Particles without Fabs were used as a negative control (black circles). Each data point represents a bound fraction value measured after approximately 2-3 h of incubation. The solid lines are guides to the eye.*

#### S4. Fitting parameters of dose-response curves in Figure 3C

*Table S2. Fitting parameters of the dose-response curves in Figure 3C.*

| Fit parameters | Series 1 | Series 2 | Series 3 | Series 4 | Series 5 |
|---|---|---|---|---|---|
| **Best-fit values** | | | | | |
| Bottom | 0 | 0 | 0 | 0 | 0 |
| Top | 1.99 | 2.01 | 1.86 | 2.44 | 3.12 |
| n | 1.03 | 1.03 | 1.06 | 0.97 | 0.90 |
| EC50 (pM) | 985 | 1007 | 868 | 1418 | 2185 |
| **95% CI** | | | | | |
| Top | 1.71 to 2.45 | 1.71 to 2.53 | 1.58 to 2.36 | 1.80 to 4.45 | 2.06 to 9.66 |
| n | 0.95 to 1.11 | 0.95 to 1.12 | 0.96 to 1.17 | 0.84 to 1.10 | 0.77 to 1.03 |
| EC50 | 743 to 1454 | 747 to 1534 | 640 to 1358 | 812 to 4210 | 1017 to 15186 |
| **Goodness of Fit** | | | | | |
| Degrees of Freedom | 67 | 67 | 67 | 67 | 67 |
| R squared | 0.99 | 0.99 | 0.99 | 0.99 | 0.99 |
| Sum of Squares | 0.04 | 0.05 | 0.07 | 0.11 | 0.12 |
| Sy.x | 0.03 | 0.03 | 0.03 | 0.04 | 0.04 |

#### S5. Signal variations for different lactoferrin concentrations in milk samples

The sensor signals shown in Figure 4B were analyzed to examine their underlying distributions (Figure S4). Multiple statistical tests were performed to determine whether the signals from each of the six samples follow a normal distribution. As shown in Table S3, four of the six samples exhibit a normal distribution in the normalized activity.

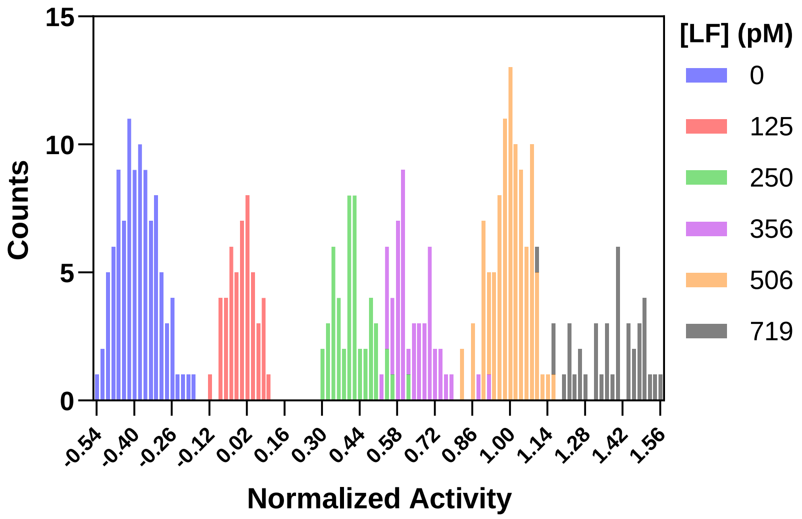


*Figure S4. Distribution of sensor signals for various lactoferrin (LF) concentrations in milk samples. Different colors correspond to the distinct lactoferrin concentrations in the samples.*

*Table S3.Test for normal distribution of the data from Figure 4B.*

| Normal distribution test | 0 pM LF | 125 pM LF | 250 pM LF | 356 pM LF | 506 pM LF | 719 pM LF |
|---|---|---|---|---|---|---|
| **D'Agostino & Pearson test** | | | | | | |
| K2 | 2.76 | 0.76 | 3.08 | 14.03 | 1.29 | 4.85 |
| P value | 0.25 | 0.68 | 0.21 | 0.00 | 0.53 | 0.09 |
| Passed normality test (alpha=0.05)? | Yes | Yes | Yes | No | Yes | Yes |
| P value summary | ns | ns | ns | - | ns | ns |
| **Anderson-Darling test** | | | | | | |
| A2* | 0.39 | 0.19 | 0.41 | 1.04 | 0.26 | 1.16 |
| P value | 0.38 | 0.90 | 0.34 | 0.01 | 0.70 | 0.00 |
| Passed normality test (alpha=0.05)? | Yes | Yes | Yes | No | Yes | No |
| P value summary | ns | ns | ns | - | ns | - |
| **Shapiro-Wilk test** | | | | | | |
| W | 0.98 | 0.98 | 0.97 | 0.91 | 0.99 | 0.93 |
| P value | 0.27 | 0.71 | 0.20 | 0.00 | 0.76 | 0.01 |
| Passed normality test (alpha=0.05)? | Yes | Yes | Yes | No | Yes | No |
| P value summary | ns | ns | ns | - | ns | - |
| **Kolmogorov-Smirnov test** | | | | | | |
| KS distance | 0.07 | 0.07 | 0.11 | 0.16 | 0.06 | 0.15 |
| P value | >0.10 | >0.10 | >0.10 | 0.00 | >0.10 | 0.01 |
| Passed normality test (alpha=0.05)? | Yes | Yes | Yes | No | Yes | No |
| P value summary | ns | ns | ns | - | ns | - |
| **Number of values** | **112** | **48** | **48** | **48** | **96** | **40** |

#### S6. Influence of Poisson noise on concentration imprecision

In Figure 4C, the concentration imprecision of different analyte concentrations was analyzed. The highest imprecision was observed at the lowest analyte concentration. This can be attributed to the Poisson nature of the binding events. Poisson noise arises because binding events occur randomly and independently, and the number of events follows a Poisson distribution. At low analyte concentrations, the frequency of binding events is lower, resulting in fewer data points and consequently, less statistical information (Figure S5A). As a result, the relative Poisson noise increases, leading to greater variability in the measured signals (Figure S5B). Additionally, a temporal effect is observed: over time, the number of switches decreases, leading to greater variability. Since concentration estimates are derived from signal measurements, greater signal imprecision directly translates to greater concentration imprecision.

The relative Poisson noise is calculated as the square root of the number of switches (Poisson noise) divided by the number of switches (signal). In Figure S5C, the coefficient of variation (CV) of the signal is plotted against the relative Poisson noise, showing similar magnitudes, indicating that Poisson noise is a large contributor to the measurement imprecision. Note that Figure S5C shows the CV of the signal, which is proportional to the CV of the concentration and scales with the inverse slope of the calibration curve.

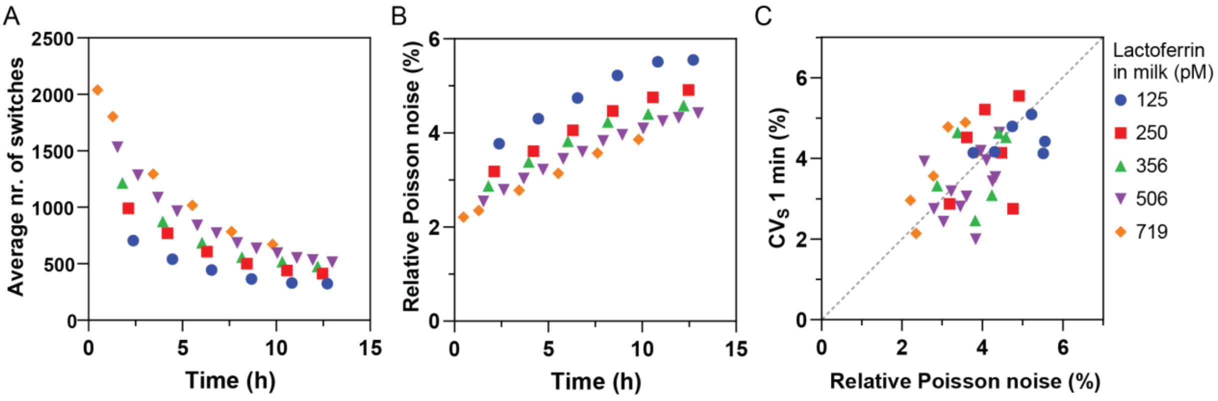


*Figure S5. Poisson noise analysis of the data in Figure 4C. (A) The average number of switches over time for each lactoferrin concentration. (B) Relative Poisson noise over time for each lactoferrin concentration. (C) Relative Poisson noise compared to the coefficient of variation of the signal of 1-min recordings ($CV_S$ 1 min). Different symbols and colors represent the different lactoferrin concentrations.*

#### S7. Dilutional linearity

The dilutional linearity of measurements on milk samples was evaluated using a range of dilution factors, as shown in Figure S6. Each sample was diluted using five different dilution factors and measured four times per dilution. To assess the consistency of the results, the experiments were repeated on different sensing cartridges. The data were fitted using log-log linear regression, yielding slope values close to -1 (Table S4). This confirms that the sensor maintains dilutional linearity for dilution factors between 500 and 8000, demonstrating that matrix effects do not significantly influence measurements within this range.

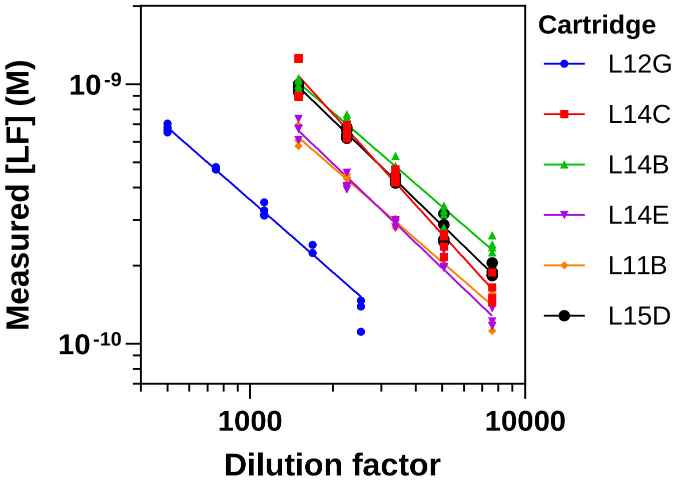


*Figure S6. Measured lactoferrin concentration as a function of dilution factor to assess dilution linearity. Each sample was diluted five times, and each dilution was measured four times on the same cartridge, with all individual data points plotted. The experiment was repeated using different cartridges, represented by different colors and symbols. The solid lines indicate log-log linear regression fits used to determine the slope. Fitting parameters are shown in Table S4.*

*Tabel S4. Fitting parameters of the log-log linear regressions in Figure S6.*

| Cartridge | **L12G** | **L14C** | **L14B** | **L14E** | **L11B** | **L15D** |
|---|---|---|---|---|---|---|
| **Best-fit values** | | | | | | |
| Y Intercept | -6.67 | -5.27 | -6.09 | -5.96 | -6.29 | -5.77 |
| Slope | -0.93 | -1.17 | -0.91 | -1.01 | -0.92 | -1.02 |
| **95% CI (profile likelihood)** | | | | | | |
| Y Intercept | -6.79 to -6.54 | -5.85 to -4.63 | -6.29 to -5.88 | -6.25 to -5.66 | -6.53 to -6.04 | -5.93 to -5.62 |
| Slope | -0.97 to -0.88 | -1.36 to -0.99 | -0.98 to -0.86 | -1.11 to -0.93 | -0.99 to -0.85 | -1.07 to -0.97 |
| **Goodness of Fit** | | | | | | |
| Degrees of Freedom | 18 | 18 | 18 | 18 | 18 | 18 |
| R squared | 0.99 | 0.94 | 0.99 | 0.98 | 0.98 | 0.99 |
| Sum of Squares | 5.21E-21 | 1.40E-19 | 2.09E-20 | 1.57E-20 | 1.16E-20 | 9.41E-21 |
| Sy.x | 1.70E-11 | 8.82E-11 | 3.40E-11 | 2.95E-11 | 2.54E-11 | 2.29E-11 |

#### S8. Concentration imprecision over time

Figure S7 shows the concentration imprecision ($CV_C$ 1 min) over time for a milk sample containing 506 pM lactoferrin, measured on 22 cartridges. An increase in imprecision is observed over time. This trend can be attributed to two main factors. First, the dynamic range of the sensor signal gradually decreases, leading to a reduced signal-to-noise ratio. As the signal becomes weaker, fluctuations in concentration measurements become more pronounced, resulting in increased imprecision in later measurements. Second, Poisson noise plays a role as fewer events are detected over time, as described in Section S6.

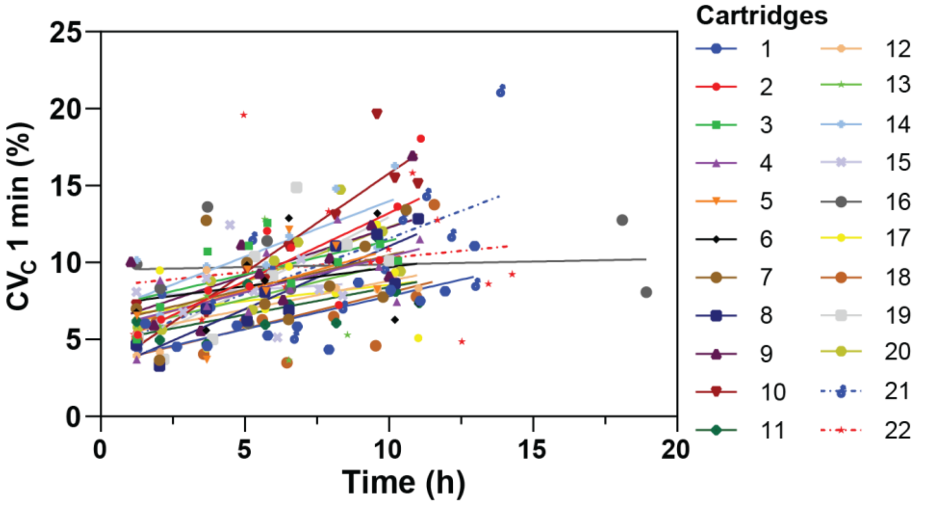


*Figure S7. Concentration imprecision ($CV_C$ 1 min) over time for a milk sample with 506 pM lactoferrin, measured on 22 cartridges. Each cartridge is represented by a unique color and symbol. The solid and dotted lines show the linear fit for each cartridge.*

#### S9. Calibration strategies compared

Three calibration strategies were compared in Figure 5: (1) repeated calibration approximately every four hours, (2) a single calibration at the start, and (3) a fixed calibration curve (Equation 1: $S_{min}$ = -0.5, $S_{max}$ = 4, n = 1, EC50 = 1.03 nM). To further validate the calibration strategies, we analyzed data from ten cartridges (Figure S8). All methods yielded similar outcomes, confirming that a fixed calibration curve is effective, demonstrating the robustness of the sensor.

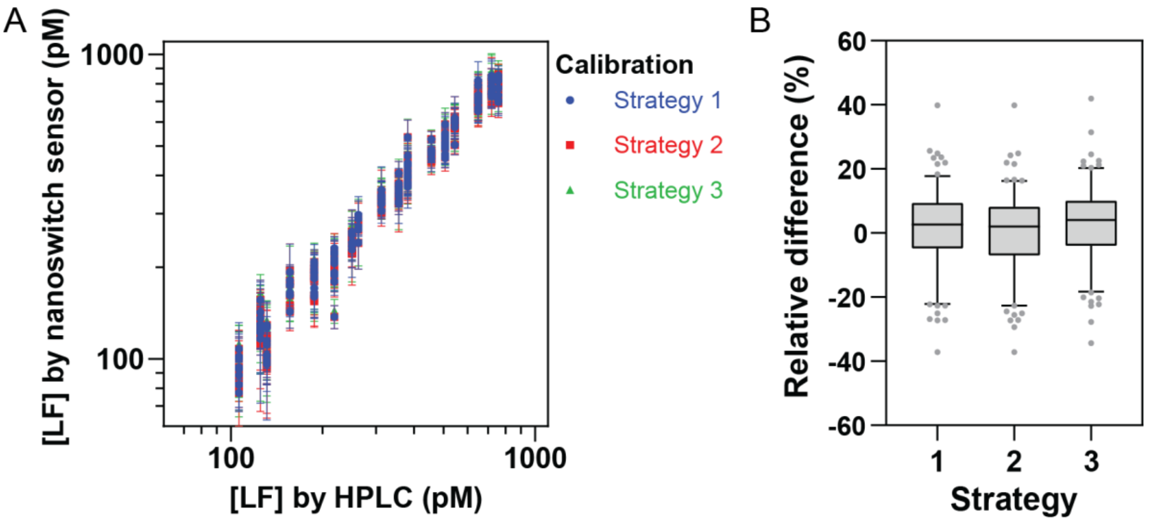


*Figure S8. Comparison of quantification strategies across all measurements from Figure 5D. (A) The correlation between lactoferrin concentrations determined by the nanoswitch sensor and HPLC is shown for Strategy 1 (blue), Strategy 2 (red), and Strategy 3 (green). (B) Relative difference of each strategy with respect to the values obtained from HPLC. The boxes illustrate the distribution of all data points: the whiskers represent the 95% range, the horizontal line within the box indicates the median, the box itself encompasses the interquartile range (50% of the data within the 95% range), and the gray dots represent outliers beyond the 95% range.*

#### S10. Relative difference between nanoswitch and HPLC measurements

Figure 5D in the body text demonstrates the correlation between nanoswitch and HPLC measurements. To further evaluate the differences between the two methods, we analyzed the relative difference from the line y = x, as shown in Figure S9. The relative difference follows a normal distribution (p = 0.125; D'Agostino & Pearson test) with a mean of 4.4% and a standard deviation of 8.5% (Figure S9B). The absolute values of the relative differences are shown in Figure S9C. The Mean Absolute Relative Difference (MARD) is 8.2% for concentrations within the calibration range (125–719 pM, Figure S9D).

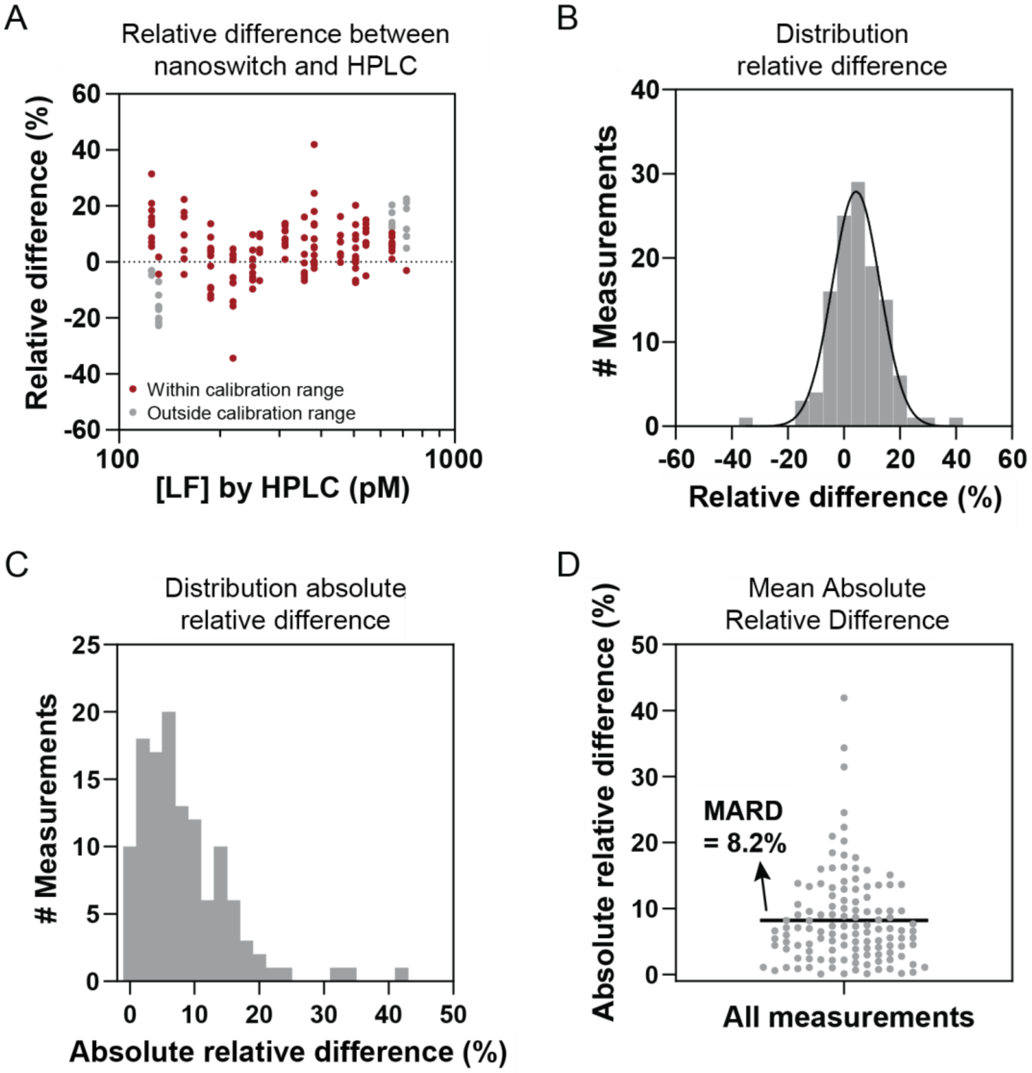


*Figure S9. Deviations between nanoswitch and HPLC concentrations. (A) Relative difference values calculated with respect to the correlation y = x. Concentration values within the calibration range are shown in red. Concentrations outside the calibration range are shown in grey and are excluded from the analysis in subsequent panels. (B) Histogram of the relative difference of all measurements, fitted with a normal distribution (p = 0.125; D'Agostino & Pearson test). The fit yields a mean relative difference of 4.4% with a standard deviation of 8.5%. (C) Distribution of the absolute relative difference of all measurements. (D) All data points are shown, with the Mean Absolute Relative Difference (MARD) represented by the black horizontal line at 8.2%.*
